# Astrocyte-derived ApoE is Required for the Maturation of Injury-induced Hippocampal Neurons and Regulates Cognitive Recovery After Traumatic Brain Injury

**DOI:** 10.1101/2021.01.13.425890

**Authors:** Tzong-Shiue Yu, Yacine Tensaouti, Elizabeth P. Stephanz, Elizabeth E. Rafikian, Mu Yang, Steven G. Kernie

## Abstract

Polymorphisms in the apolipoprotein E (ApoE) gene confer a major genetic risk for the development of late-onset Alzheimer’s disease (AD) and are predictive of outcome following traumatic brain injury (TBI). Alterations in adult hippocampal neurogenesis have long been associated with both the development of AD and recovery following TBI, and ApoE is known to play a role in this process. In order to determine how ApoE might influence hippocampal injury-induced neurogenesis, we developed a novel conditional system whereby functional ApoE from astrocytes was ablated just prior to injury. While successfully ablating 90% of astrocytic ApoE just prior to a closed cortical impact injury in mice, we observed an attenuation in the development of newly born neurons using a GFP-expressing retrovirus, but not in existing hippocampal neurons visualized with a Golgi stain. Intriguingly, animals with a “double-hit”, i.e. injury and ApoE conditionally inactivated in astrocytes, demonstrated the most pronounced impairments in the hippocampal-dependent Morris water maze test, failing to exhibit spatial memory after both acquisition and reversal training trials. In comparison, conditional knockout mice without injury displayed impairments but only in the reversal phase of the test, suggesting accumulative effects of astrocytic ApoE deficiency and traumatic brain injury on AD-like phenotypes. Together, these findings demonstrate that astrocytic ApoE is required for functional injury-induced neurogenesis following traumatic brain injury.

**Significance Statement:** ApoE has long been implicated in the development of Alzheimer’s disease and recovery from traumatic brain injury via unknown mechanisms. Using a novel conditional ablation model of mouse ApoE and subsequent tracing of individual hippocampal neurons, we demonstrate its requirement in injury-induced neurogenesis for proper dendritic arborization and cognitive function in hippocampal-dependent learning and memory tasks.

## Introduction

The presence of specific human Apolipoprotein E (ApoE) isoforms is the best known risk factor for developing late-onset Alzheimer’s disease (LOAD), although the mechanism underlying its role is unknown. In humans, polymorphic ApoE protein is derived from three alleles, E2, E3, and E4. Approximately one-quarter of all individuals are E4 carriers, and 65-80% of LOAD patients have at least one copy of the E4 allele (Mahley and Huang, 2012). Despite its clinical relevance, the function of ApoE in the brain remains largely unknown, but is believed to be critical for repairing and remodeling lipid membranes, organelle biogenesis, and neuronal dendritogenesis because of its role in cholesterol and lipid transport and metabolism (Mahley and Huang, 2012; Flowers and Rebeck, 2020).

We have recently demonstrated that systemic and developmental deficiency of ApoE results in a simplification of dendritic tree structure from newborn neurons in the dentate gyrus during both normal development and following controlled cortical impact injury (CCI). In addition, similar phenotypes were observed in humanized ApoE4 replacement mice (Tensaouti et al., 2018; Tensaouti et al., 2020). Moreover, the lack of ApoE, or the presence of human ApoE4, resulted in lower spine density, specifically in newborn neurons in the dentate gyrus. Thus, we demonstrated that ApoE is a prerequisite for both proper formation of dendritic trees and spine density of adult hippocampal newborn neurons in naïve and injured brains.

In the brain, although ApoE is observed in several types of cells, including astrocytes, microglia, and stressed neurons; the primary source of ApoE is astrocytes (Xu et al., 2006). ApoE released from astrocytes forms HDL-like lipoprotein particles with cholesterol and phospholipids (Yamazaki et al., 2019). The ApoE-containing lipoprotein particles are then taken up by cells through the LDL receptor family, including LDLR and LDLR-related protein 1 (LRP1) for ApoE-mediated lipid metabolism (Yamazaki et al., 2019). In addition, ApoE-containing lipoprotein particles bind with neuronal amyloid β (Aβ), which then undergoes endocytosis by astrocytes facilitating Aβ clearance (Liao et al., 2016).

To further illustrate the critical roles of astrocytic ApoE in lipid metabolism, neuronal plasticity, and repair, we generated ApoE conditional mice to spatially and temporally regulate ApoE expression in astrocytes using Aldh1l1-cre/ERT transgenic mice (Srinivasan et al., 2016). In the current study, we employed our novel ApoE conditional mice to specifically explore the role of astrocyte-derived ApoE in maintaining dendritic structure in both pre-existing and newly generated dentate gyrus neurons. In addition, we determined the behavioral outcomes of astrocytic ApoE deficiency and controlled cortical impact (CCI) injury ---- a well-established experimental model of traumatic brain injury (TBI) and a clinically relevant environmental insult commonly associated with increased risk of developing AD (Johnson et al., 2010; Ramos-Cejudo et al., 2018). Our results reveal that postnatal ablation of astrocytic ApoE resulted in less complexity in the dendritic tree of granular neurons, primarily in newborn dentate gyrus neurons. These observed deficits were more evident in brains from ApoE conditionally deficient mice following CCI. Most remarkably, these morphologic changes appear to be functionally relevant, as demonstrated by the pronounced spatial memory deficits in ApoE conditionally deficient mice with CCI, in both the acquisition and reversal phases of the Morris water maze test. In uninjured mice, the lack of astrocytic ApoE did not impair acquisition memory, but impaired reversal memory, indicating additive effects of astrocytic ApoE deficiency and CCI, consistent with our findings at the neuronal level. Hence, our findings suggest that astrocytic ApoE is critical for the development and function of newly born granular neurons in the dentate gyrus, and this effect is particularly evident in the injured brain.

## Material and methods

### Animals

All procedures in this research were performed following the animal protocol approved by the Institutional Animal Care and Use Committee at Columbia University. Subject mice were housed humanely and cared for inside the barrier facility managed by the Institute of Comparative Medicine at Columbia University. Food and water were provided ad libitum for all mice, and were kept on a 12 h light/dark cycle with lights on at 7 A.M. B6; FVB-Tg(Aldh1l1-cre/ERT2)1Khakh/J and Ai14(RCL-tdT)-D lines were ordered from The Jackson Laboratory to allow expression of tdTomato in astrocytes specifically in a tamoxifen-induced manner.

The generation of the apoef/f mouse was conducted by Knockout Mouse Program (KOMP) at University of California, Davis (project number 456). Briefly, exon 1 and exon 2 of murine ApoE were targeted and flanked with loxp sequences; therefore, the start of transcription is impaired. Upon generation of germline transmitted apoe f/f mice, they were mated with Tg(Pgk1-flpo)10Sykr mice (Stock No. 011065, The Jackson Laboratory) to remove the neo cassette. These progeny were mated with Aldh1l1-creERT2 and Ai14 to get Aldh1l1-cre/ERT2; Ai14; apoef/f, littermates with other genotypes were used as controls. To identify the genotype of apoef/f mice, primer pairs (forward: 5’-CCGTGCTGTTGGTCACATTGC-3’; reversed: 5’-GCATGCACTGTCTTGTATCCTATGTAG −3’) were used to perform PCR. The band with size ~500 bps represents wild-type apoe gene and 750 bps PCR fragment indicates the loxp flanked apoe gene.

### Controlled cortical impact (CCI) and retroviral injections

Six-week old mice were used for the experiments. For controlled cortical impact injury (CCI), we used a standard protocol with a controlled cortical impact device (Leica Impact One, Leica Biosystems) to generate brain injuries as described (Tensaouti et al., 2020). Following anesthesia with isoflurane, mice were placed in a stereotactic frame. A midline incision was made, the soft tissues were reflected, and a 5X5 mm craniotomy was made between bregma and lambda 1 mm lateral to the midline. The injury was generated with a 3 mm stainless steel tipped impact device with deformation of 0.7 mm, a constant speed of 4.4 m/s, and a duration of 300 milliseconds. Mice operated on but not impacted were used as sham groups.

Immediately following the injury (ctr, sham (Ctr-Sham)=5; cko, sham (cKO-Sham)=5; ctr, CCI (Ctr-CCI)=5; cko, CCI (cKO-CCI)=8), 1 microliter of retrovirus with enhanced green fluorescent protein (eGFP) expressing vector (RV-CAG-eGFP, 1*10^8 transducing unit/mL, Salk Institute) was infused stereotactically into the dentate gyrus at an injection rate of 0.1 μl per minute by a micro infusion pump (KD Scientific) via a 10 μl microsyringe (Model 801, Hamilton). The following coordinates were used: Anterior/Posterior: −2.0 mm, Medial/Lateral: +/-1.55 mm and Dorsal/Ventral: −2.0 mm in reference to bregma. After surgery, the scalp was closed with sutures and mice were placed in their cages and allowed to recover from anesthesia.

### Tissue processing and immunohistochemistry

Four weeks after surgery, mice were perfused with 4% paraformaldehyde (PFA, 441244, Sigma-Aldrich) and brains were harvested for post-fixation in 4% PFA overnight. The brains were then sectioned serially using a vibratome (VT1000S, Leica) with thickness of 50 um. All sections encompassing the hippocampus were harvested sequentially into 6-well plates.

A set of sections was washed with 1xPBS then permeabilized with 0.3% of Triton X-100/1xPBS (PBST) at room temperature. Sections were blocked with normal donkey serum (5% NDS, 017-000-121, Jackson ImmunoResearch). The primary antibodies to label ApoE (1:5000, 7333, ProSci) and eGFP (1:500, A11122, Invitrogen) were added and incubated with sections overnight at room temperature with 0.02% of sodium azide. The following day, sections were washed with PBST, then biotinylated donkey anti-goat IgG was used to label ApoE primary antibody and Alexa 488-conjugated donkey anti-rabbit IgG eGFP antibody (1:200, both from Jackson ImmunoResearch) at 4°C overnight. On the third day, sections were washed again with PBST, then Alexa 647-conjnugated streptavidin was incubated with sections for visualization of apoE at room temperature for three hours. Then sections were washed with 1xPBS before mounted on the slides with Vectashield mounting medium (H-1500, Vector Laboratories).

#### Quantification of ApoE-expressing cells

To verify if ApoE expression was suppressed in the brains because of tamoxifen treatment in a cre-dependent manner, ApoE-expressing cells were counted using unbiased stereological method. Samples (Ctr-Sham=3; cKO-Sham=3; Ctr-CCI=3; cKO-CCI=3) were analyzed under a Zeiss microscope (Axio Imager M2, Zeiss). The whole hippocampus was traced under a 10x objective lens. The sample grids (500*500 mm) were determined randomly and cells within the counting frames (75*75 mm) were counted under a 40X objective. To prevent artifacts that resulted from sectioning, a dissector height of 30 mm was used. The ApoE-expressing cell number was estimated using the built-in weighted section thickness method with a coefficient of error less than 0.1 (Stereology Investigator, MBF).

#### Golgi-Cox staining

Standard Golgi-Cox staining method was followed (Zaqout and Kaindl, 2016). Stock solutions of 5% (w/v) of potassium dichromate (P5271, Sigma), mercuric chloride (AC201430250, Fisher Scientific), and potassium chromate (216615, Sigma) were prepared and stored at room temperature in the dark. The working solution was prepared by mixing 50 mL of potassium dichromate with mercuric chloride stock solution, then 40 mL of potassium chromate stock solution was added followed by 100 mL of distilled water. The prepared working solution was left at room temperature covered with foil for at least 48 hours to allow for precipitate formation.

Four weeks after surgery (Ctr-Sham=4; cKO-Sham=4; Ctr-CCI=3; cKO-CCI=4), mice were sacrificed and brains were removed fresh and washed with distilled water. The washed brains were then transferred to a small bottle that contained 10 mL of clear Golgi-Cox working solution and stored at room temperature. One day later, the solution was renewed and left for 7 days for impregnation. After impregnation, brains were transferred to new bottles containing 30% sucrose solution and store at 4°C and protected from light. One day later, the brains were transferred into new bottles with sucrose solution for 4-7 days at 4°C in the dark.

Brains were cut using a vibratome (VT1000S, Leica) and serial 100 um sections encompassing the entire hippocampus were harvested. Sections were positioned on gelatin-coated slides to allow further development with 3:1 ammonia solution (1054261000, Milipore Sigma) for 8 minutes. After development, tissue was dehydrated using serial ethanol solutions. Sections were then kept in Xylene (23400, Electron Microscopy Sciences) before mounting using permount (SP15-500, Fisher Scientific).

#### Neuronal morphological analysis

Sections with either eGFP-expressing cells or Golgi-Cox stained cells were visualized using a Zeiss microscope (Axio Imager M2, Zeiss) with a Hamamatsu camera (Orca-R2, Hamamatsu). Stack images with 1 mm interval on z-axis were acquired (Stereology Investigator, MBF) under a 20x objective lens. Neurolucida 360 (MBF) was used to reconstruct and analyze the dendritic structure. Total length of dendrites, nodes of dendrites, and Sholl analysis were performed.

### Behavioral Tests

Four weeks after surgery, subject mice were acclimated to a vivarium room dedicated for housing behavioral subjects for a week and then a carefully selected test battery was conducted in this order: elevated plus-maze, open field, Morris water maze acquisition, Morris water maze reversal, and cued and contextual fear conditioning. Inter-test intervals were 3 to 7 days. All behavioral experiments were done between 10am and 4pm during the light phase. The experimenter was blind to genotype and injury information while conducting the experiments and subsequent data analysis. For Ctr-Sham group, eight males and six females were included; Ctr-CCI group, eight males and seven females; cKO-Sham, five males and seven females; and seven males and eight females in cKO-CCI group.

#### Elevated plus-maze

The elevated plus-maze consists of two open arms (30cm x 5cm) and two closed arms (30 x 5 x 15 cm) extending from a central area (5 x 5 cm). Photo beams embedded at arm entrances register movements. Room illumination was approximately 5 lux. The test began by placing the subject mouse in the center, facing a closed arm. The mouse was allowed to freely explore the maze for 5 min. Time spent in the open arms and closed arms, the junction, and number of entries into the open arms and closed arms, were automatically scored by the MED-PC V 64bit Software (Med Associates). At the end of the test, the mouse was gently removed from the maze and returned to its home cage. The maze was cleaned with 70% ethanol and wiped dry between subjects.

#### Open Field Test

The Open Field is the most commonly used test for spontaneous exploratory activity in a novel environment, incorporating measurements of locomotion and anxiety-like behaviors. The Open Field test was performed following previously described protocols (Yang et al., 2012). Exploration was monitored during a 60 min session with Activity Monitor Version 7 tracking software (Med Associates Inc.). Briefly, each mouse was gently placed in the center of a clear Plexiglas arena (27.31 x 27.31 x 20.32 cm, Med Associates ENV-510) lit with dim light (~5 lux), and was allowed to ambulate freely. Infrared (IR) beams embedded along the X, Y, Z axes of the arena automatically track distance moved, horizontal movement, vertical movement, stereotypies, and time spent in center zone. Data were analyzed in six, 5-min time bins. Areas were cleaned with 70% ethanol and thoroughly dried between trials.

#### Morris water maze acquisition and reversal

Spatial learning and memory were assessed in the Morris water maze following previously described protocols (refs). No visible trials were run before or after hidden platform trials in the current study. The 122cm circular pool was filled 45 cm deep with tap water and rendered opaque with the addition of nontoxic white paint (Crayola). Water temperature was maintained at 23°C±1. The proximal cue was one sticker taped on the inner surface of the pool, approximately 20 cm above the water surface. Trials were videotaped and scored with Ethovision XT 12 (Noldus). Acquisition training consisted of four trials a day for seven days. Each training trial began by lowering the mouse into the water close to the pool edge, in a quadrant that was either right of, left of, or opposite to, the target quadrant containing the platform (12 cm in diameter). The start location for each trial was alternated in a semi-random order for each mouse. The hidden platform remained in the same quadrant for all trials during acquisition training for a given mouse, but varied across subject mice. Mice were allowed a maximum of 60 s to reach the platform. A mouse that failed to reach the platform in 60 s was guided to the platform by the experimenter. Mice were left on the platform for approximately 15 s before being removed. After each trial, the subject was placed in a cage lined with absorbent paper towels and allowed to rest under an infrared heating lamp for 1 min. Two hours after the completion of the last training trial, the platform was removed and mice were tested in a 60 s probe trial. Parameters recorded during training days were latency to reach the platform, total distance traveled, and swim speed. Time spent in each quadrant and number of crossings over the trained platform location and over analogous locations in the other quadrants were used to analyze probe trial performance. All trials were recorded and analyzed with the Ethovision XT video tracking software (Noldus Information Technology Inc).

#### Fear Conditioning

Fear conditioning was assessed following previously described protocols (Yang et al., 2012). Training and conditioning tests are conducted in two identical chambers (Med Associates, E. Fairfield, VT) that were calibrated to deliver identical foot shocks. Each chamber was 30 cm × 24 cm × 21 cm with a clear polycarbonate front wall, two stainless side walls, and a white opaque back wall. The bottom of the chamber consisted of a removable grid floor with a waste pan underneath. When placed in the chamber, the grid floor connected with a circuit board for delivery of scrambled electric shock. Each conditioning chamber was placed inside a sound-attenuating environmental chamber (Med Associates). A camera mounted on the front door of the environmental chamber recorded test sessions which were later scored automatically, using the VideoFreeze software (Med Associates, E. Fairfield, VT). For the training session, each chamber was illuminated with a white house light. An olfactory cue was added by dabbing a drop of imitation lemon flavoring solution on the metal tray beneath the grid floor. The mouse is placed in the test chamber and allowed to explore freely for 2 min. A pure tone (5kHz, 80 dB) which serves as the conditioned stimulus (CS) was played for 30 s. During the last 2 s of the tone, a foot shock (0.5 mA) was delivered as the unconditioned stimulus (US). Each mouse received three CS-US pairings, separated by 90 s intervals. After the last CS-US pairing, the mouse was left in the chamber for another 120 s, during which freezing behavior is scored by the VideoFreeze software. The mouse was then returned to its home cage. Contextual conditioning is tested 24 h later in the same chamber, with the same illumination and olfactory cue present but without foot shock. Each mouse was placed in the chamber for 5 min, in the absence of CS and US, during which freezing is scored. The mouse was then returned to its home cage. Cued conditioning is conducted 48 h after training. Contextual cues were altered by covering the grid floor with a smooth white plastic sheet, inserting a piece of black plastic sheet bent to form a vaulted ceiling, using near infrared light instead of white light, and dabbing vanilla instead of lemon odor on the floor. The session consisted of a 3 min free exploration period followed by 3 min of the identical CS tone (5kHz, 80dB). Freezing was scored during both 3 min segments. The mouse was then returned to its home cage. The chamber was thoroughly cleaned of odors between sessions. % freezing on Day 1 was analyzed to indicate the immediate reaction to receiving foot shocks, % freezing on Day 2 and Day 3 was analyzed to reflect contextual conditioning and cued conditioning, respectively.

### Statistical analysis

All statistical analyses excluding behavioral tests were performed using Prizm (ver. 9, GraphPad). Results are presented as mean ± SEM. A value of p less than 0.05 was considered statistically significant. A One-way ANOVA with Bonferroni post-hoc analysis was used to determine the statistical difference in ApoE-expressing cell number among the experimental groups. To compare the total length of dendrites and nodes, unpaired t-test was used. The data from Sholl analyses were analyzed using Two-way ANOVA with Bonferroni post-hoc analysis.

For behavioral results, statistical analyses and graphs were generated in SigmaPlot (Systat Software, Inc.). All data is presented as mean ± SEM with a p-value < 0.05 considered statistically significant. WT and cKO were analyzed separately. A Two-way RM ANOVA, with time and operation (sham vs. CCI) as the two factors, was performed on distance traveled, center time, and vertical movement data collected from the open field test. For Morris Water Maze data, latency to platform, swim distance, and swim speed were all analyzed using Two-way RM ANOVA with Day as the within-subject factor and operation (sham vs. CCI) as the between-subject factor. Mean % time spent in each quadrant for each group during the probe trial was analyzed using One-way RM ANOVA, followed by Dunnett’s post-hoc test, to see if the animals spent significantly more time in the trained quadrant than in the other three quadrants while platform crossings during the probe trials were analyzed using an unpaired t-test. Percentage of time spent freezing during the contextual and cued phases of Fear Conditioning was analyzed using an unpaired t-test.

## Results

### Generation of ApoE conditional knockout animal and experimental design

In order to bypass the developmental effects of conventional ApoE-deficient and humanized ApoE4 mice, we developed a conditional knockout model of ApoE using a Cre-loxp strategy. ApoE^*f/f*^ mice were derived using a construct in which the first and second exons of the murine ApoE gene was flanked with loxP sites (*Materials and Methods, Fig. 1A*). To verify that mice had germline transmission of ApoE^*f/f*^, primers targeting exon 2 of the ApoE gene and upstream loxP site were used to create DNA fragments by performing traditional PCR. The 750-bps fragment indicated the ApoE^*f/f*^ mouse whereas the 500-bps fragment the ApoE^*wt/wt*^ mouse, and the 500/750 fragments were found in heterozygous ApoE^*f/wt*^ mice (*Fig. 1B*).

**Figure 1.**
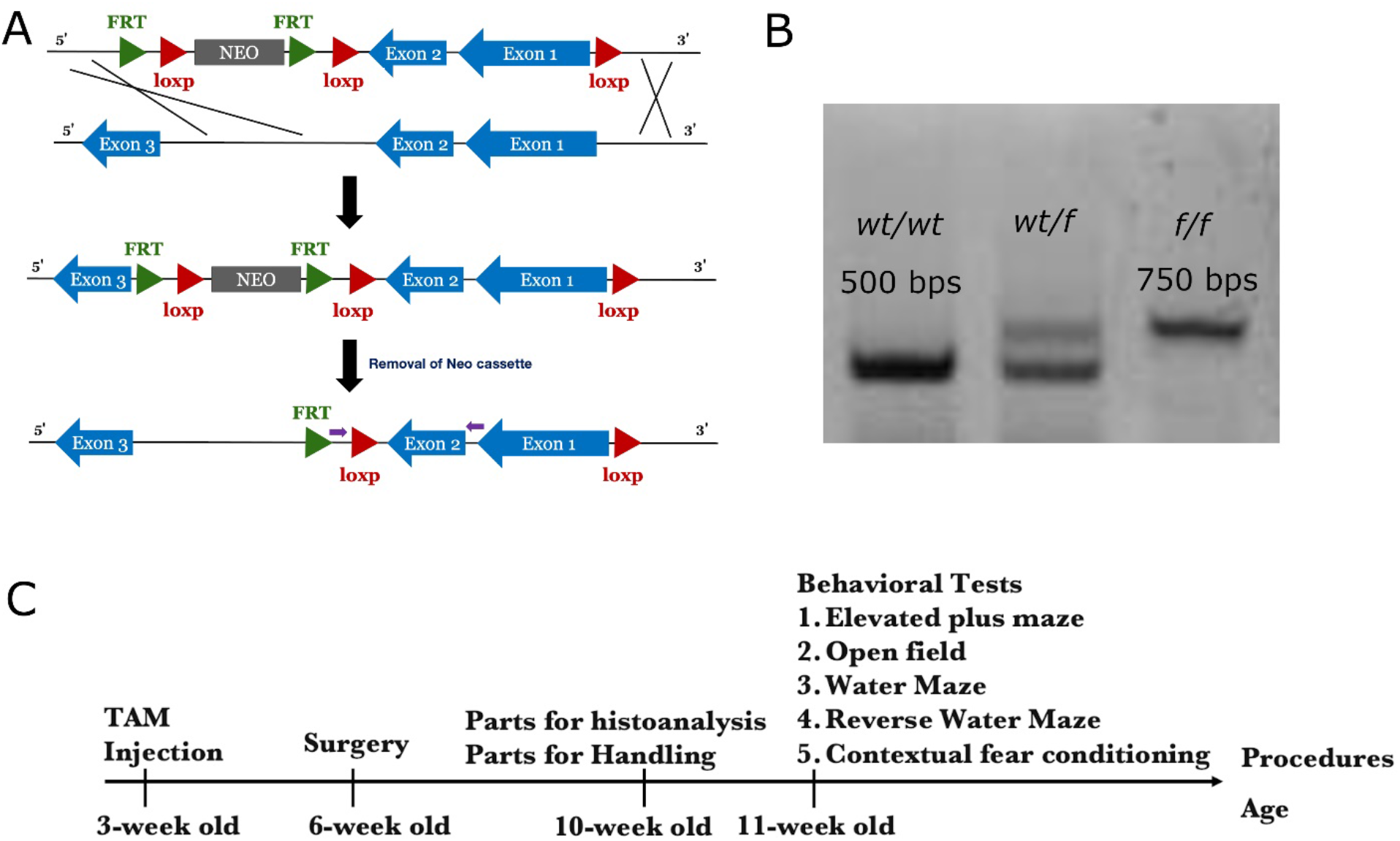
Overview of the generation of ApoE conditional knockout animals and experimental outline. (A) To generate ApoE conditional knockout mice, exon 1 and exon 2 of the ApoE gene were flanked with loxp sites to allow for cre-dependent recombination. The primers for genotyping were designed to flank exon 2 of the ApoE gene with or without one of the loxp sites. (B) The 500 bp DNA fragment generated by PCR indicates mice without the loxp flanked ApoE, the 750 bp result indicates mice with two copies of loxp-flanked ApoE, and the hemizygous mice generate one 500 bp and one 750 bp fragment. (C) Timeline of experiments. Mice received tamoxifen at three weeks of age. At six weeks of age, sham or CCI surgery was performed. A subset of mice received eGFP-expressing retrovirus targeted to the dentate gyrus at the conclusion of surgery. Four weeks after viral infection, mice were perfused to analyze the morphology of newborn neurons or stained using Golgi-Cox method. Habituation was performed on the remaining mice. Following habituation, the elevated plus maze, open field, Morris water maze and contextual fear conditioning tasks were performed.

ApoE expression has been observed in astrocytes, microglia, and stressed neurons in the brain, though astrocytes are the primary source (Xu et al., 2006; Mahley, 2016; Kockx et al., 2018). To investigate the specific role of astrocytic ApoE in the brain, we used Aldh1l1-cre/ERT transgenic mice with validated specificity for cre/ERT-dependent recombination in astrocytes (Srinivasan et al., 2016). Aldh1l1-cre/ERT transgenic mice were mated with ApoE^*f/f*^ mice to allow temporally specified deletion of the ApoE gene in astrocytes, and tdTomato-expressing reporter mice Ai14 were used for visualization. At three weeks of age, ApoE^*f/f*^ and littermate control mice received one injection of tamoxifen each day for 3-5 days, to trigger deletion of ApoE in astrocytes (*Fig. 1C*). When the treated mice were six weeks old, they received either a sham operation or CCI; and a subset of them received an intracranial injection of eGFP-expressing retrovirus to label dividing newborn neurons in the dentate gyrus. They were perfused four weeks after surgery to analyze the morphology of eGFP-expressing mature newborn neurons (*Fig. 1C*). For mice not infected with the retrovirus, brains were harvested four weeks after surgery for Golgi-Cox staining to determine the dendritic complexity of neurons in the dentate gyrus (*Fig. 1C*). Finally, for behavioral testing, habituation started four weeks after surgery and was performed once a day for five days. Following habituation, elevated plus maze and open field tasks were performed to determine whether the lack of astrocytic ApoE affected stress and anxiety. To investigate if the reduction of astrocytic ApoE impaired learning and memory, Morris water maze and contextual fear conditioning tasks were performed (*Fig. 1C*).

### Aldh1l1-cre/ERT induced knockout of ApoE in astrocytes

To investigate the role of astrocytic ApoE in neuronal plasticity and maturation of newborn neurons in the dentate gyrus in both naïve and injured brains, ApoE conditional knockout mice were crossed with Aldh1l1-cre/ERT BAC transgenic mice to induce astrocyte-specific cre-dependent gene recombination to allow ablation of ApoE expression in astrocytes in a tamoxifen-inducible manner (Srinivasan et al., 2016). The mice were also crossed with Ai14, to verify the efficiency of cre-mediated recombination by allowing tdTomato expression where this occurs. Mice received one injection of tamoxifen daily for 3-5 days. To determine the efficiency of ApoE deletion in astrocytes in both naïve and injured mice, tamoxifen-treated mice received a sham operation or CCI at six weeks of age (*Ctr-Sham=3; Ctr-CCI=3; cKO-Sham=3; cKO-CCI=3; Fig. 1A*). Four weeks after surgery, brain sections were collected for ApoE staining. Most of the ApoE-expressing cells in the hippocampus expressed tdTomato-from sham-operated and injured non-ApoE^*f/f*^ siblings, and the morphology of these cells confirmed that they were astrocytes (*Fig. 2A-C, arrows*). As compared to controls, the number of ApoE-expressing cells in the hippocampus in the ApoE^*f/f*^ conditional knockout mice was dramatically reduced (*Fig. 2D-F, arrows*). Using unbiased stereology, 1~1.5*10^6 ApoE-expressing cells in the hippocampus were counted in both naïve and injured controls (*Fig. 2J*). In the Aldh1l1-cre/ERT; ApoEf/f mice, the number of ApoE-expressing cells was significantly reduced to 0.1~0.5*10^6 cells (*Fig. 2J, One-way ANOVA, F=13.13, p=0.0019; Bonferroni post-hoc test; *: p<0.05, **: p<0.01*). Therefore, Aldh1l1-cre/ERT induced knockout of ApoE in approximately 90% of astrocytes in both sham-operated and injured mice. In the absence of the Aldh1l1-cre/ERT transgene, the injection of tamoxifen did not result in the recombination of loxP-flanked genes as no tdTomato-expressing cells were observed and ApoE expression was preserved (*Fig. 2G-I*). Hence, Aldh1l1-cre/ERT triggers efficient knockout of the ApoE gene in astrocytes in a tamoxifen-dependent manner.

**Figure 2.**
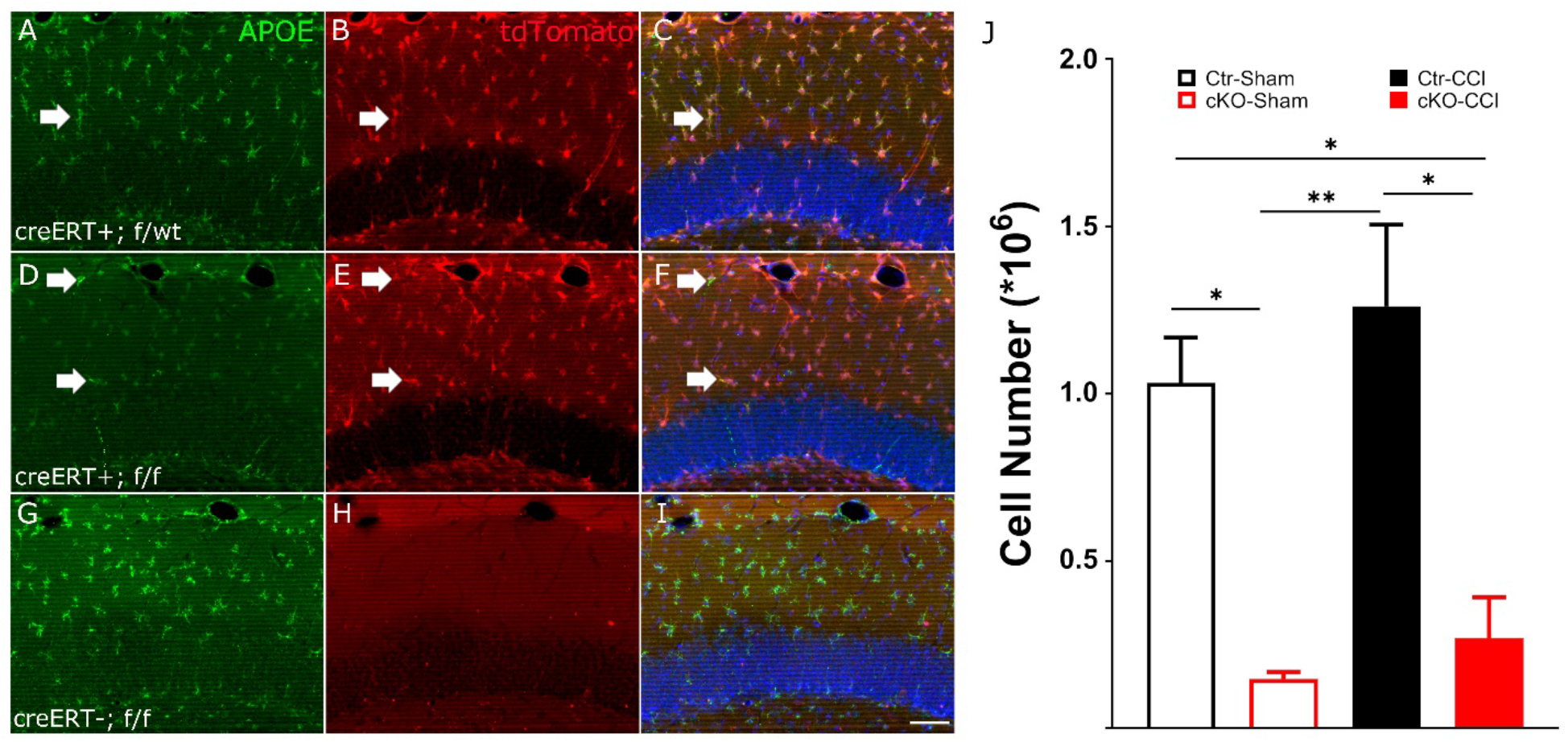
Tamoxifen and aldh1l1-creERT dependent knockdown of ApoE expression in astrocytes. (A-C) 7 weeks after tamoxifen in control littermates, ApoE expression was observed in most aldh1l1-dependent tdTomato-expressing cells in the hippocampus, as shown by arrows. (D-F) In apoe^f/f^ mice, ApoE expression in tdTomato-expressing cells is dramatically reduced. (G-I) In the absence of aldh1l1-creERT, the expression of ApoE in the hippocampus remained unchanged in mice receiving tamoxifen and the expression of cre-dependent tdTomato was not observed. This indicates that no spontaneous recombination of loxp-flanked DNA fragments ocurred. (J) Significant reduction in the number of ApoE-expressing cells in the hippocampus in aldh1l1-creERT; apoe^f/f^ mice in both sham-operated and CCI mice. Data are presented as means ± SEM. Scale bar in I = 50 μm.

### Reduction of astrocytic ApoE mildly impairs the dendritic complexity of pre-existing neurons in the dentate gyrus

As a relay hub, the dentate gyrus receives information from the entorhinal cortex and transmits it back via CA3 and CA1. Adult neurogenesis in the dentate gyrus and the connections between CA3 and the dentate gyrus are known to play a critical role in encoding new memories and distinguishing them from existing ones to prevent memory overlap (Miller and Sahay, 2019). The abundant expression of ApoE in the molecular layer of the dentate gyrus where the dendrites of granular neurons reside suggest that ApoE is critical in dendritic formation and maintenance. To elucidate the potential role of astrocytic ApoE in maintaining the dendritic trees of existing neurons in the dentate gyrus in sham-operated and injured mice, Golgi-Cox staining method was deployed four weeks after surgery to select random neurons for analysis (*Fig. 3A, C, E and G. Ctr-Sham=5 mice, 137 neurons; Ctr-CCI=4 mice, 372 neurons; cKO-Sham=4 mice, 100 neurons; cKO-CCI=4 mice, 260 neurons. Scale Bar=100 μm*). The dendritic trees of stained neurons were then restructured for morphological analysis using Neurolucida 360 (*Fig. 3B, D, F, and H*). Using Sholl analysis, the distribution of dendrites in the dentate gyrus in sham-operated mice lacking astrocytic ApoE differed with respect to distance from the soma when compared with controls (*Fig. 3I. Two-way ANOVA with Bonfferoni post-hoc analysis. Experimental Groups: p<0.0001, F_1,7050_=27.02; Radius: p<0.0001, F_29,7050_=414.3; Interaction: p=0.8601, F_29,7050_=0.7229*). Additionally, there was no difference observed in the total length of the dendrites and nodes (*Fig. 3J and K. Two-tail unpaired t-test for both, total length: p=0.1997; node: p=0.7230*). Therefore, dendritic trees of existing dentate neurons lacking astrocytic ApoE demonstrated slightly less complexity than neurons in sham-operated controls. Similarly, Sholl analysis revealed that the lack of astrocytic ApoE did not impair the distribution of dendrites of the dentate neurons (*Fig. 3L. Two-way ANOVA with Bonferroni post-hoc analysis. Experimental Groups: p=0.1096, F_1,18900_=2.561; Radius: p<0.0001, F_29,18900_=703.3; Interaction: p=0.9982, F_29,18900_=0.4037*) or the total length of the dendrites and branching points in the injured mice, as compared to sham controls (*Fig. 3M and N. Two-tail unpaired t-test for both, total length: p=0.6711; node: p=0.1967)*. Hence, the reduction of astrocytic ApoE modestly impairs dendritic complexity and maintenance of dendrites in the pre-existing dentate neurons in the injured or control brain.

**Figure 3.**
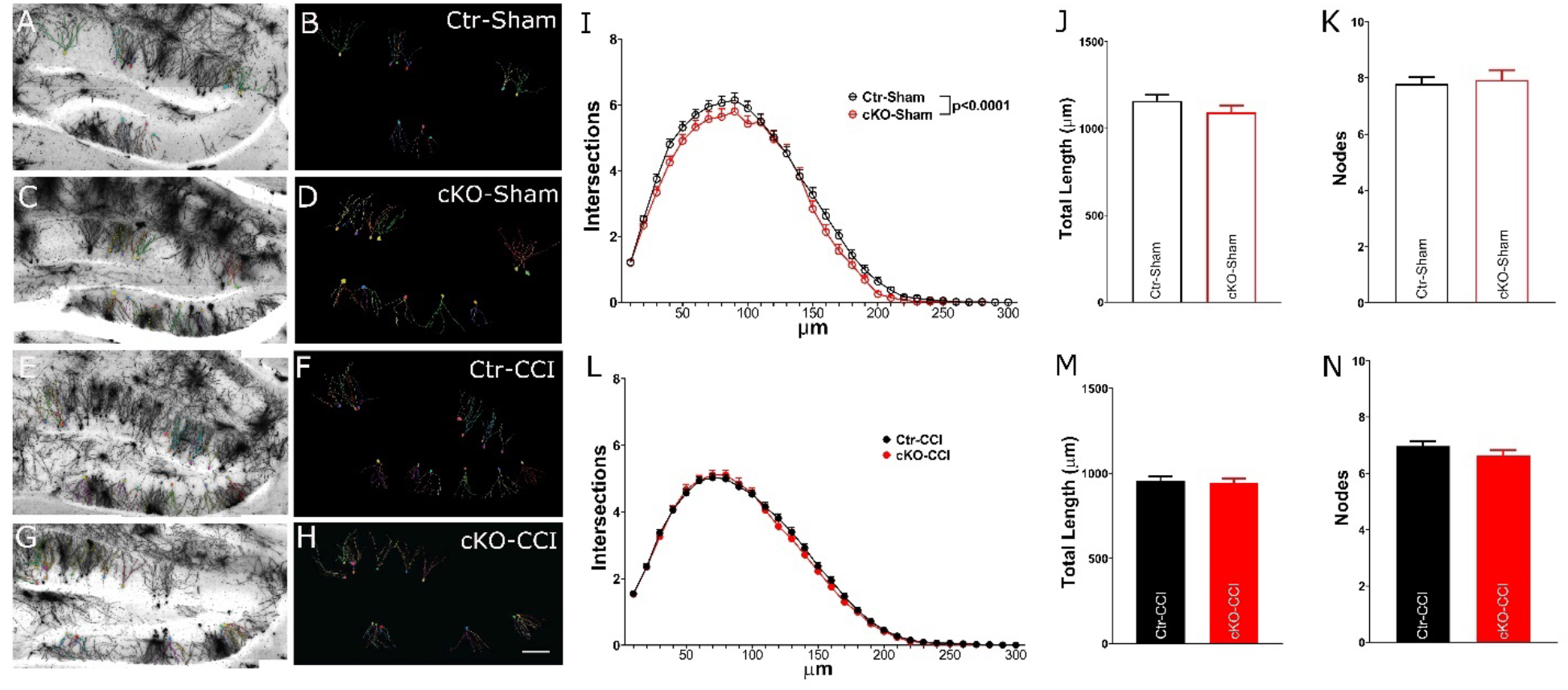
The morphology of Golgi-Cox stained neurons in the hippocampus mice without astrocytic ApoE. (A-H) The left panels demonstrate Golgi-Cox stained neurons in the dentate gyrus and the the right panel depicts individual analyzed neurons. (I-K) In sham-operated mice, two-way ANOVA analysis revealed a significant difference in Sholl analysis between the experimental groups (I) but there was no difference was observed either in the total length (J) or branching nodes (K) of the dendrites in mice lacking astrocytic ApoE compared with control littermates. (L-N) No significant difference was observed in dendritic complexity, total length, and branching nodes in the dentate gyrus in the injured mice with or without astrocytic ApoE. Data are presented as means ± SEM. Scale bar in H = 100 μm.

### Depletion of astrocytic ApoE markedly impairs dendritic complexity of newly born neurons in injured mice

In order to determine whether ablation of ApoE impairs the development of newly born dentate gyrus neurons, both controls and conditional knockout mice received an eGFP-expressing retrovirus to label dividing newborn neurons at the time of surgery (*Fig. 4A. Ctr-Sham=4 mice, 212 neurons; Ctr-CCI=5 mice, 133 neurons; cKO-Sham=5 mice, 197 neurons; cKO-CCI=4 mice, 214 neurons. Scale Bar=100 μm.*). Analysis took place four weeks later to allow for infected newborn neurons to mature. In sham-operated groups, Sholl analysis demonstrated slightly increased complexity of dendritic trees in the ApoE conditional knockout mice (*Fig 4A, B and E. Two-way ANOVA with Bonferroni post-hoc analysis. Experimental Groups: p<0.0001, F_1,11396_=17.67; Radius: p<0.0001, F_27,11396_=391.8; Interaction: p=0.3264, F_27,11396_=1.101)* and a Bonferroni *post-hoc* analysis revealed that this slight increase in complexity occurred proximal to the soma specifically (*\* p=0.0483*). However, the difference was not observed in the total length and node number (*Fig 4F and G. two-tail unpaired t-test for both, total length: p=0.2423; node: p=0.1519*). In injured mice, deletion of astrocytic ApoE resulted in less sophisticated dendritic distribution and the post-hoc Bonferroni test revealed a reduction in complexity in the distal dendrites (*Fig. 4C, D and H. Two-way ANOVA with Bonferroni post-hoc analysis. Experimental Groups: p<0.0001, F_1,9660_=40; Radius: p<0.0001, F_27,9660_=250.1; Interaction: p=0.0001, F_27,9660_=2.320*). The lack of astrocytic ApoE in the injured mice also resulted in a decrease in the total length of dendrites in the injured group (*Fig. 4I and J. Two-tail unpaired t-test for both, total length: p=0.0456; node: p=0.5348*). Thus, we conclude that astrocytic ApoE is critical for the development of dendritic arborizations of newborn neurons in injured mice.

**Figure 4.**
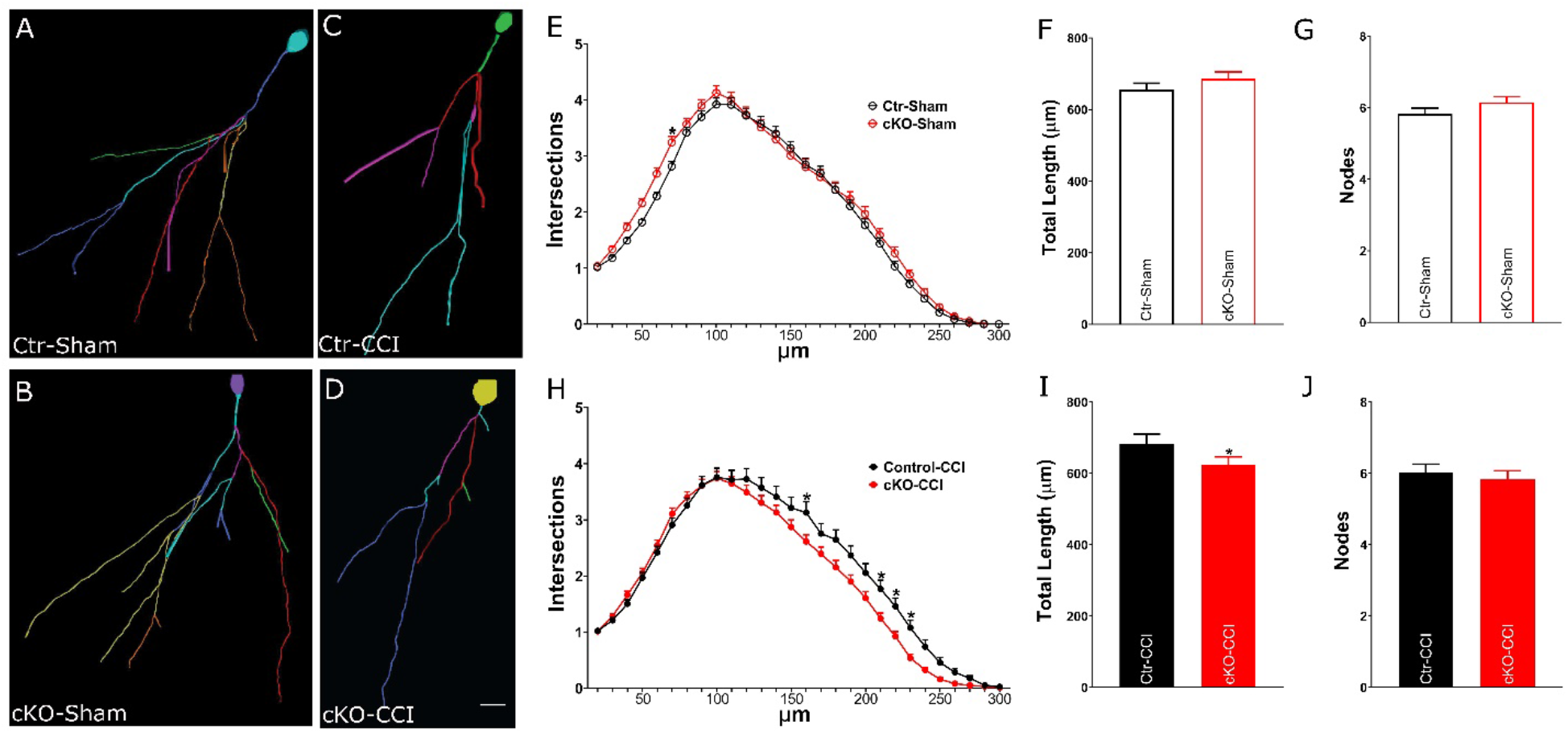
The reduction in astrocytic ApoE impairs dendritic complexity in newborn neurons in the injured dentate gyrus. (A-D) Reconstructed dendritic trees of newborn neurons 4 weeks after being injected with a GFP-expressing retrovirus in the dentate gyrus. (E-G) Sholl analysis demonstrates that in sham-operated mice, the reduction of astrocytic ApoE results in an increased proximal dendritic complexity of newborn neurons (E). However, this difference was not observed in the total length (F) and branching nodes (G). (H-J) In injured mice, lack of astrocytic ApoE impairs distal dendritic complexity of newborn neurons (H) and total dendritic length (I) but not in branching nodes (J). Data are presented as means ± SEM. *:p<0.05. Scale bar in D = 10 μm.

### Spatial memory was impaired in ApoE cKO mice and worsened by CCI

In the Morris water maze test, CCI did not impair spatial memory in control (Ctr) mice, either in acquisition or reversal (Fig. 5A-B). Using One-Way RM ANOVA followed by Dunnett’s test for post-hoc comparisons, we found that both Ctr-Sham and Ctr-CCI mice spent significantly more time in the trained quadrant than in the other three quadrants in the probe trial following acquisition training (the main effect of “quadrant”: Ctr-Sham, F_3,13_=31.55, p<0.01; Ctr-CCI, F_3,14_=8.22, p<.01). Post-hoc comparisons indicate that both Ctr-Sham and Ctr-CCI spent significantly more time in the trained quadrant than in the other three quadrants (p<.01 for all comparisons). Similarly, in the probe trial following reversal training, both Ctr-Sham and Ctr-CCI mice spent significantly more time in the trained quadrant (the main effect of “quadrant”: Ctr-Sham, F_3,13_=7.06, p<0.01; Ctr-CCI, F_3,14_=4.69, p<.01). Post-hoc comparisons indicated that the differences between the trained quadrant and all other three quadrants were significant for Ctr-Sham (p<.05 for all comparisons). For Ctr-Sham, the differences between the trained quadrant and the quadrants opposite of, and right to, the trained quadrants were significant (p<.01 for both comparisons).

**Figure 5.**
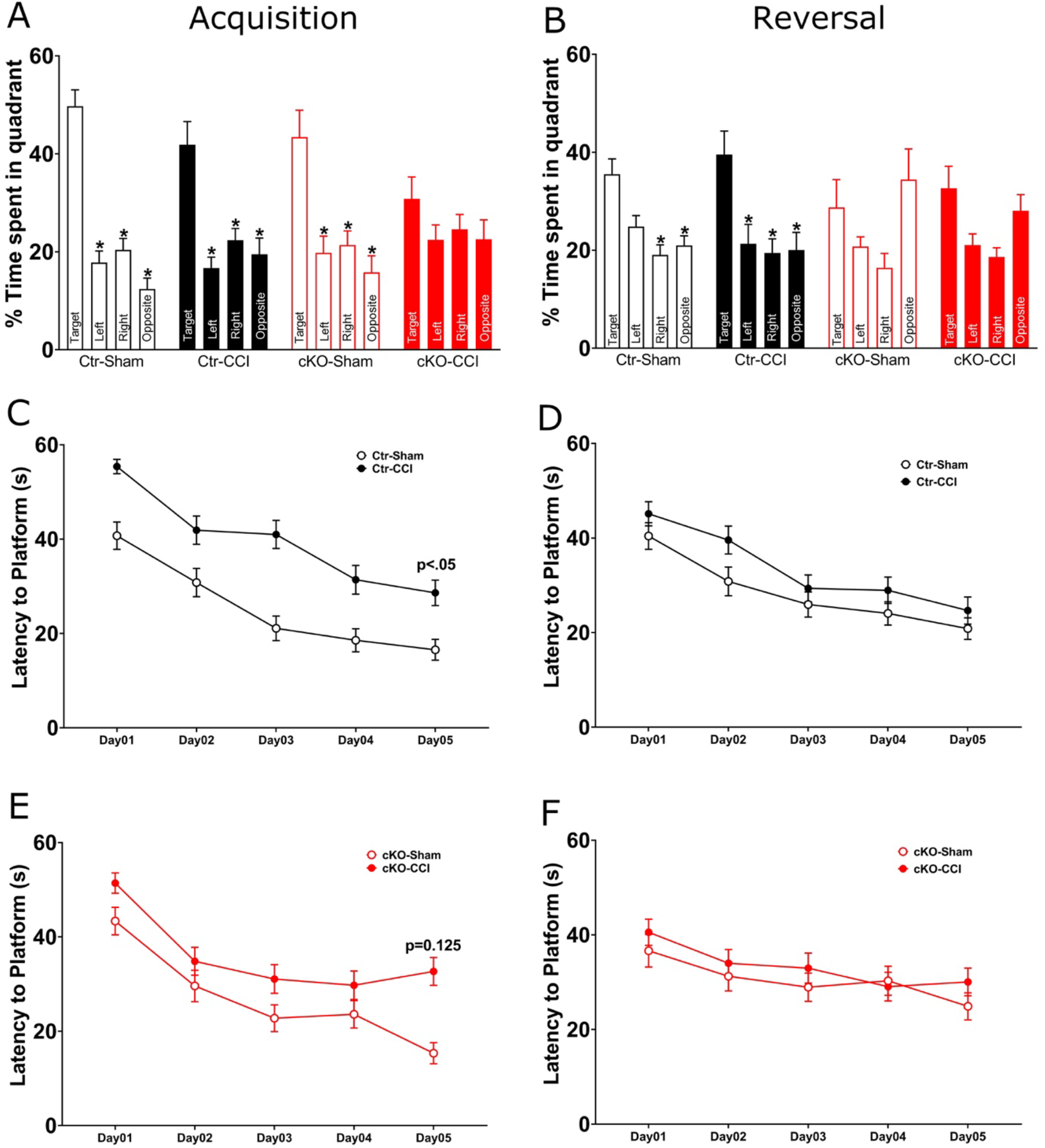
Impaired spatial memory in astrocytic ApoE deficient mice was worsened by CCI in the Morris water maze test. (A) Astrocytic ApoE deficiency did not affect spatial memory in uninjured mice after acquisition training, but did impair memory in cKO mice with CCI. (B) Astrocytic ApoE deficiency alone was sufficient to impaired reversal memory. The same impairment was seen in cKO mice with or without CCI. (C, E). During the acquisition training session, injured mice exhibited longer latency to platform than sham-operated mice. This was significant in controls and approaching significance in cKO mice. *p<.05 versus the trained quadrant. Data are presented as means ± SEM.

In ApoE conditional knockout mice (cKO), an additive effect of ApoE deficiency and CCI was seen, with uninjured cKO mice only exhibiting impaired memory following acquisition, and injured cKO mice exhibiting impaired memory following both acquisition and reversal. While cKO-Sham mice exhibited a clear preference for the trained quadrant in the probe trial following acquisition (F_3,13_=7.53, p<.01), spending significantly more time in the trained quadrant than in the other three quadrants (p<.01 for all comparisons), this group failed to exhibit quadrant preference in the probe trial following reversal training (F_3,13_=2.32, NS), indicating modest impairment. A more pronounced impairment was seen in cKO mice with CCI, which did not exhibit a preference for the trained quadrant in the probe trials after acquisition training (F_3,14_=0.85, NS) or the reversal training (F_3,14_=3.09, p<.05; posthoc comparisons did not reveal significant differences between the trained quadrant and any of the other three quadrants).

Latency to platform, swim distance, and swim speed were analyzed by genotype (Ctr and cKO) (Fig 5C-F). Two-way RM ANOVA was used, with “Day” as the within subject factor, and operation (sham vs. CCI) as the between subject factors. During acquisition training days, latency to platform was significantly longer in Ctr-CCI mice than in Ctr-Sham mice (F_1,27_=10.00, p<.01). No significant effect of CCI was found for latency to platform in cKO mice (F_1,25_=2.53, NS). During reversal training days, no significant effects of CCI were found on latency to platform in Ctr mice (F_1,27_=1.22, NS) or cKO groups (F_1,25_=0.23, NS).

The increase in latency in the Ctr-CCI group is probably related to a decrease in swim speed in this group (Fig 6A-H). Compared to Ctr-Sham mice, swim speed was significantly slower in Ctr-CCI mice during acquisition (F_1,27_=4.47, p<.05) and reversal (F_1,27_=9.75, p<.01). In cKO mice, swim speed was slower in cKO-CCI mice during acquisition (F_1,25_=5.59, p<.05). No speed difference was found in reversal (F_1,25_=1.17, NS). In Ctr mice, swim distance was shorter in Ctr-CCI mice than in Ctr-Sham mice during acquisition (F_1,27_=9.05, p<.01) but not during reversal (F_1,27_=0.05, NS). In cKO mice, swim distance was not different between the Sham and CCI group during acquisition (F_1,25_=0.39, NS) or reversal (F_1,25_=0.23, NS). In sum, Ctr mice exhibited normal learning and memory in the water maze acquisition and reversal, despite CCI’s effect in reducing swim speed. In contrast, a partial deficit was found in cKO-Sham mice, which exhibited normal acquisition memory but impaired reversal memory. This deficit was exacerbated by CCI, as evidenced by cKO-CCI mice showing impaired memory after both acquisition and reversal learning trials.

**Figure 6.**
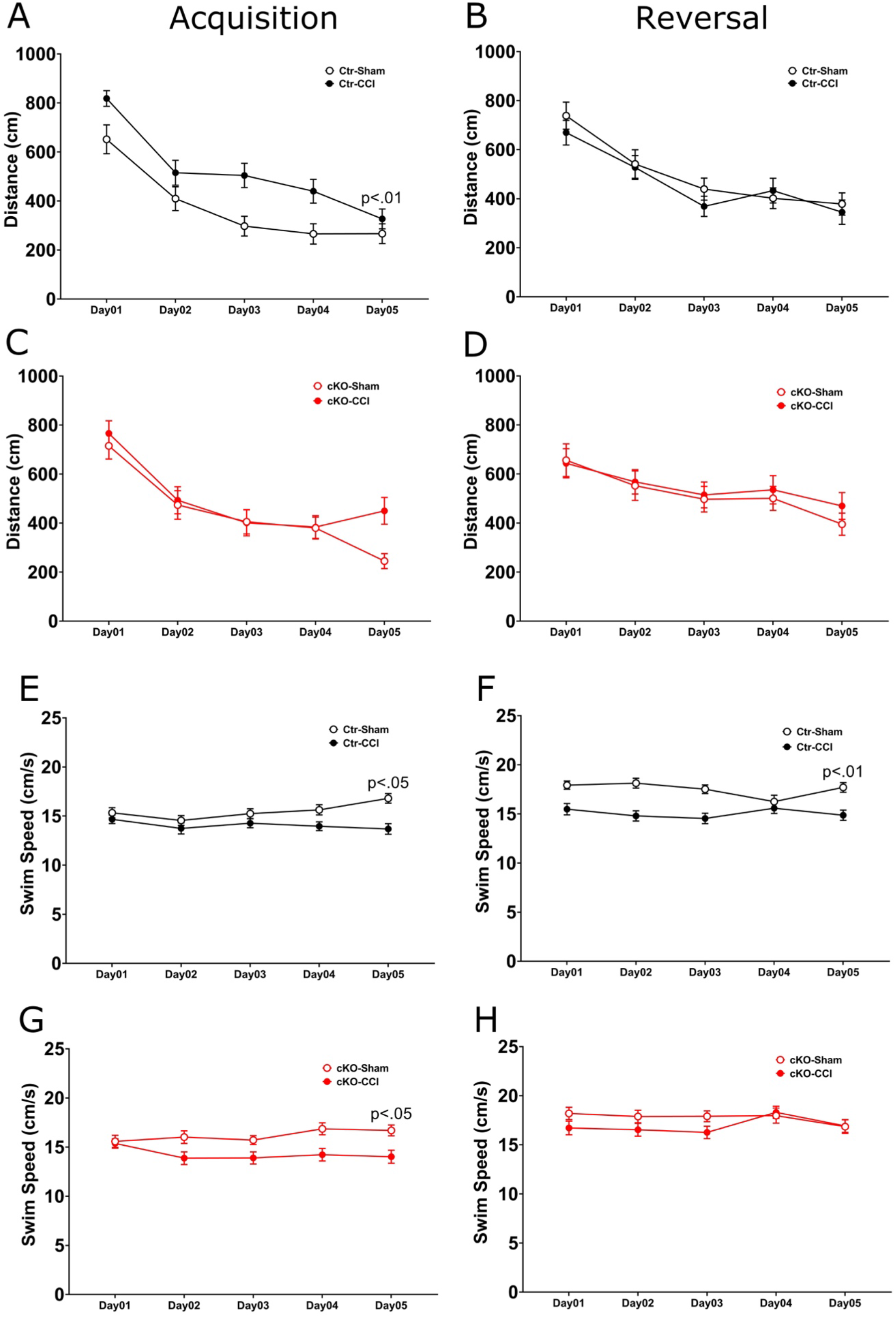
Water maze speed and distance was modestly affected by CCI but not ApoE deficiency. (A, C, E, G). During acquisition, injured mice swam a longer distance to reach the hidden platform as compared to the control groups, however, there was no CCI effect on swim distance in mice lacking astrocytic ApoE. The injury also resulted in slower swimming speed despite the presence of astrocytic ApoE. (E, F, F, G). In reversal learning, all mice moved a similar distance to reach the new position of the platform. Interestingly, the swimming speed was slower in the Ctr-CCI group. However, there was no difference observed between cKO-Sham and cKO-CCI.

In the fear conditioning test (Fig 7,D), no significant differences were found in post-training freezing. In the context conditioning test on Day 2, a significant operation effect was found (F_1,52_=5.14, p<.05) with CCI mice showing less freezing. Genotype (F_1,52_=0.68, NS) and genotype x operation interaction (F_1,52_=0.28, NS) were not significant. No significant differences were found in pre-cue freezing or cued freezing on Day 3.

**Figure 7.**
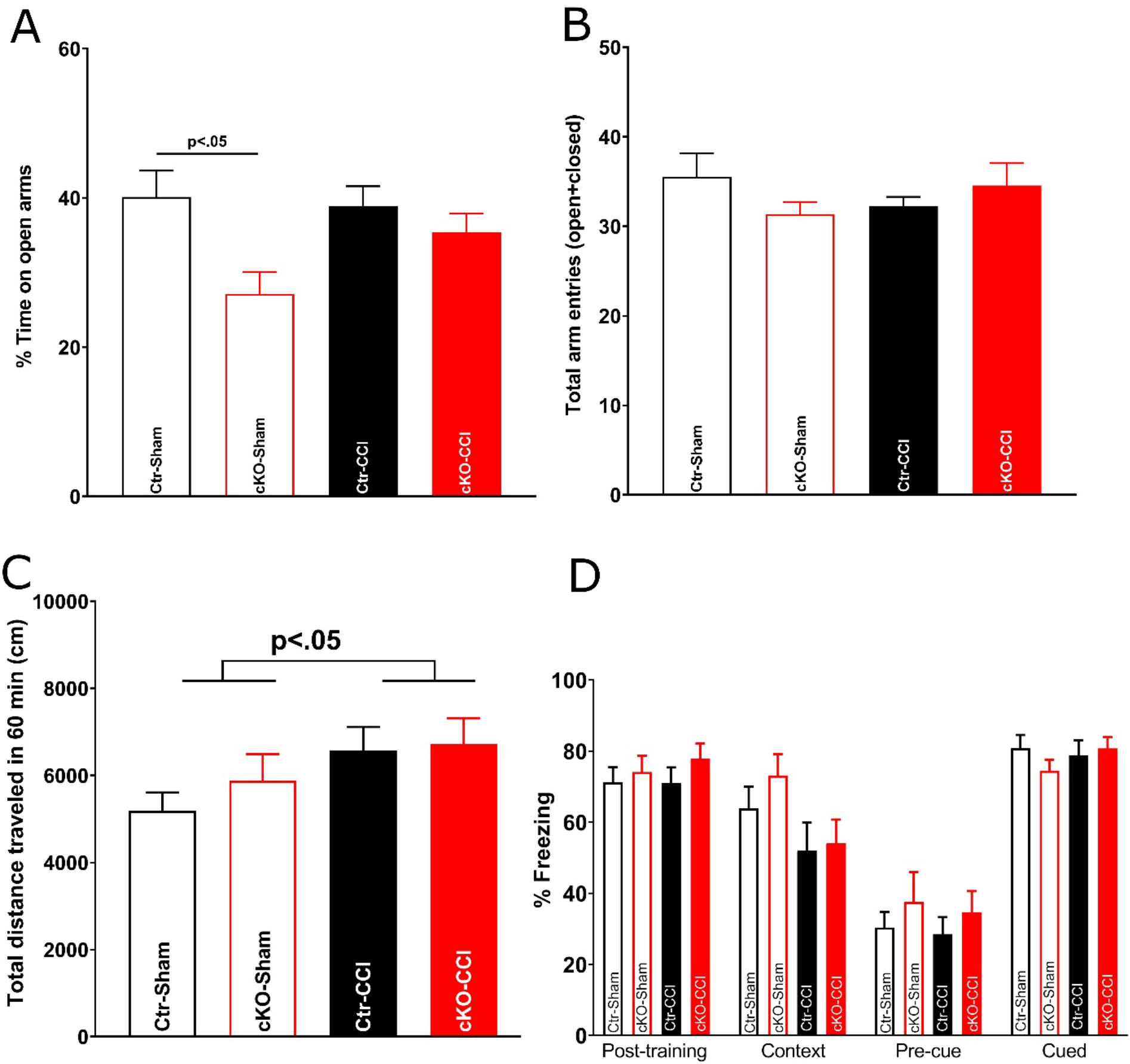
Elevated plus maze, open field, and fear conditioning. (A, B), In the elevated plus maze task, the cKO-Sham mice spent significantly less time in the open arm compared with Ctr-Sham. No difference was observed between Ctr-CCI and cKO-CCI. For the total entries of arms, no difference was observed among four groups. (C). In the open field test, injured mice did travel significantly longer distance compared with sham-operated mice, indicating CCI does not impair mobility. (D) In the contextual fear conditioning, no difference was observed in post-training task. In day 2 context-dependent test, the injured mice resulted in less time freezing compared with Sham groups. In day 3 cue-dependent test, no difference was observed among four groups.

### Anxiety-like behavior and exploratory activity

As illustrated in Fig 7 A and B, in the elevated plusmaze test, two-way ANOVA revealed a significant effect of genotype on % time in the open arms, which is reduced in cKO mice (F_1,48_=7.88, p<.01). The main effect of operation (F_1,48_=1.48, NS) and the interaction between genotype and operation (F_1,48_=2.59, p=0.11) were not significant. For total arm entries, no significant effects were found for genotype (F_1,48_=0.22, NS), operation (F_1,48_=0.00, NS), or the genotype x operation interaction F_1,48_=2.56, p=.11). In the open field test (Fig. 7C), two-way ANOVA revealed a significant effect of operation, with CCI mice showing increased distance traveled in the open field (F_1,52_=4.11, p<.05). Effects of genotype (F_1,52_=0.59, NS) and genotype x operation interaction F_1,52_=0.25, NS) were not significant. Results of the open field test ruled out gross motor impairments in cKO mice.

## Discussion

ApoE is abundant in the brain and is primarily synthesized by astrocytes, yet its role in neuronal development and maintenance remains surprisingly unclear. To determine how astrocytic ApoE might affect adult neurogenesis, we generated conditional ApoE knockout mice and crossed them with well-established astrocyte-specific Aldh1l1-creERT mice to ablate the expression of ApoE in astrocytes (Srinivasan et al., 2016). After tamoxifen treatment, we observed an approximately 90% reduction of ApoE in the hippocampus. This decrease in astrocytic ApoE resulted in mild impairments in the complexity of the existing dendritic tree and more pronounced ones in newborn granular cells. This effect was most evident in injury-induced neurons in the dentate gyrus. This attenuation in granular cell complexity may underlie impairments in reversal learning observed in the Morris water maze. Though mice learned to find the hidden platform during the training session, ApoE-deficient mice failed to retrieve the platform’s location in the reversal learning task. A controlled cortical impact model of traumatic brain injury augmented the deficits observed in the sham groups. Thus, the lack of astrocytic ApoE simplified the structure and shortened the total length of the dendritic trees of newborn neurons in the dentate gyrus and impaired acquisition and reversal memories.

ApoE is ubiquitously expressed in the brain. Using mice with the ApoE locus replaced by eGFP, ApoE expression was seen in 75% of astrocytes, 10% of microglia, and essentially no neurons (Xu et al., 2006). However, additional studies have demonstrated ApoE in pericytes, oligodendrocytes, choroid plexus and adult neural progenitors (Pitas et al., 1987; Bruinsma et al., 2010; Yang et al., 2011; Nelissen et al., 2012; Achariyar et al., 2016; Flowers and Rebeck, 2020). In addition to astrocytes, the adult neural progenitor population has also been shown to express Aldh1l1 (Foo and Dougherty, 2013). To distinguish its function in the present study, tdTomato-expressing granular cells in the dentate gyrus were only occasionally observed using Ai14 mice to trace the Aldh1l1 active cells via expression of tdTomato. Thus, we achieved efficient and specific ablation of ApoE expression in astrocytes with minimal deletion in the stem cells themselves, where ApoE has been shown to be expressed and required for progenitor proliferation (Yang et al., 2011).

We utilized the Golgi-Cox staining method to investigate whether astrocytic ApoE affected the existing dendritic structure in the dentate gyrus. In the sham-injured groups, a small but statistically significant difference in the dendritic structure was observed using Sholl analysis between controls and astrocyte-specific ApoE knockout mice. In previous studies using constitutive ApoE-deficient mice and MAP2 as a marker of hippocampal dendritic trees, lack of ApoE resulted in a decrease of neuropil in the inner molecular layer of the dentate gyrus starting from 4-months of age (Masliah et al., 1997). However, using the same approach, others reported that no significant difference was observed in the dentate gyrus (Anderson et al., 1998). Furthermore, conflicting results were reported using a transgenic approach to overexpress human ApoE driven by cell-specific promoters or using humanized ApoE mice (Masliah et al., 1995; Dumanis et al., 2009; Jain et al., 2013). Using human ApoE4 targeted replacement mice, dendritic trees were less complex compared with wild-type mice, although the structure in the dentate gyrus was not studied (Dumanis et al., 2009). When using a cell-specific promoter to drive the expression of ApoE, overexpression of human ApoE4 in neurons resulted in few dendrites in CA1 neurons, but no deficit was observed when overexpressed in astrocytes (Jain et al., 2013). The caveats in these published reports include poor control of the expression level of ApoE when using a transgenic approach and the developmental effects when using constitutively ApoE-deficient mice.

The novelty here is the development and use of a conditional knockout of ApoE to suppress its expression in a tissue-specific and temporally controlled manner. Seven weeks after beginning repression of ApoE expression in astrocytes, we observed modestly less complex dendritic trees in dentate gyrus granular neurons. Unlike what was observed in the sham-operated groups, we did not observe similar deficits in dendritic arborizations in injured brains. ApoE is known to facilitate cholesterol transport from astrocytes to neurons. Thus, ApoE controls some aspects of dendritic development and remodeling in an LDL-dependent manner (Pitas et al., 1987; Mamotte et al., 1999; Mahley, 2016; Flowers and Rebeck, 2020). Therefore, the lack of ApoE may be implicated in the impairments observed in the remodeling of dendritic trees. Using humanized ApoE4 transgenic mice, ApoE appears to regulate the thickness of the molecular layer of the dentate gyrus thirty days post-surgery (Champagne et al., 2005). This observation suggests that the expression of ApoE4, which appears to function in some circumstances as a dominant-negative form of ApoE, worsened the impairments caused by acquired brain injury. Thus, although the lack of astrocytic ApoE resulted in further deficits caused by brain injury, it will be critical to investigate the long-term effects in the injured brain caused by the lack of astrocytic ApoE (Tensaouti et al., 2018).

We previously demonstrated the impact of ApoE in newborn neurons in the dentate gyrus and uncovered its required role in the development of dendritic trees of newborn neurons. In ApoE-deficient mice, newborn neurons showed less complex dendritic trees when compared with wild-type mice (Tensaouti et al., 2018). Adult neurogenesis in the dentate gyrus is facilitated by astrocytes, microglia, and neurons, and ApoE is synthesized in astrocytes and microglia. By specifically deleting ApoE in astrocytes after the majority of brain development has occurred, we observed more intersections of the dendritic trees in cKO-Sham mice compared with Ctr-Sham. One possibility is that newborn neurons form more complex dendritic trees that compensate for the existing neurons with less complicated dendritic trees which lack astrocytic ApoE. There may also be compensation occurring that is derived from ApoE in microglia or adult neural progenitors. Consistent with the previous study, newborn neurons in injured mice grew less complicated dendritic trees when lacking ApoE (Tensaouti et al., 2020). Thus, astrocytic ApoE is critical in the growth of dendrites in injury-induced newborn neurons in the dentate gyrus. It is also known that traumatic brain injury predisposes individuals towards Alzheimer’s disease and other neurodegenerative diseases later in life (Jellinger, 2004; Freire, 2012; Mahley and Huang, 2012; Johnson et al., 2017). Interestingly, impaired neurogenesis is known to accelerate the progression of neurodegenerative diseases (Hollands et al., 2017; Polis and Samson, 2021). The present study suggests that astrocytic ApoE may play a role in the progression of neurodegenerative disease resulting from impaired hippocampal neurogenesis.

To specifically investigate whether alterations in dentate gyrus neurogenesis due to lack of ApoE results in behavioral deficits, a reversal learning of the Morris water maze was performed (Vorhees and Williams, 2006; Garthe and Kempermann, 2013). Despite the lack of astrocytic ApoE, mice exhibited intact spatial memory after acquisition training. In comparison, injured mice learned more slowly compared to sham-operated mice. In the memory test after acquisition training, sham-operated mice spent more time in the target quadrant regardless of whether astrocytic ApoE was present or not. In the memory test after reversal training, astrocytic ApoE-deficient mice exhibited impaired spatial memory. A more pronounced impairment was seen in mice with double insults: absence of astrocytic ApoE and CCI. This group exhibit impaired spatial memory after both acquisition and reversal. While we did observe mildly reduced swim speed in CCI groups, it is unlikely that this reflects gross motor impairments, which could have confounded the water maze results. In the open field test, CCI groups actually transversed greater distance than uninjured mice, indicating that CCI did not impair mobility.

The Morris water maze is a well-established behavioral task that critically depends on the hippocampus to succeed in spatial learning and memory in laboratory rodents. For the reference memory version of the Morris water maze task, experimental animals learn to find the hidden platform according to the surrounding environmental cues in the task room for task acquisition. The success of acquisition is later defined by allowing the animal to swim freely in the water tank without the platform and measuring the spent time in the target area. By lesioning specific brain areas, it has been shown that an intact hippocampus is critical for accomplishing this spatial task (Vorhees and Williams, 2006; Garthe and Kempermann, 2013). Moreover, various brain injuries, including TBI, that result in a damaged hippocampus due to secondary injury also cause different levels of impairments in the Morris water maze task in a hippocampal neurogenesis-dependent manner (Blaiss et al., 2011). The present study demonstrates a correlation between the simplified dendritic trees of injury-generated newborn neurons and the poor performance in the Morris water maze where astrocytic ApoE has been deleted. Furthermore, the reversal learning of the Morris water maze revealed that without astrocytic ApoE, both sham-operated and injured mice failed to remember the new location of the platform and spend significant time in the previous target area.

Overall, it appears that adult-generated dentate gyrus neurons require ApoE not only for proper dendritic structural maturity, but also to regulate certain aspects of critical memory formation. This novel requirement for ApoE is particularly evident in the setting of traumatic brain injury and suggests the importance of future investigation particularly regarding the cell-specific function of human isoforms of ApoE that have long been correlated with impaired outcomes following TBI through unknown mechanisms. By linking ApoE function with hippocampal-specific recovery following TBI, this present work underlies the importance of additional investigation regarding how ApoE status in humans may influence both recovery and potential treatment pathways for those with both neurodegenerative and trauma-induced brain disease.

## Acknowledgements

This research was supported by National Institutes of Health/National Institute of Neurological Disorders and Stroke Grants R56-NS-089523 (SGK) and R01-NS-095803 (SGK) and the Paul Allen Foundation (SGK)

